# Beyond members of the Flaviviridae family, sofosbuvir also inhibits chikungunya virus replication

**DOI:** 10.1101/360305

**Authors:** André C. Ferreira, Patrícia A. Reis, Caroline S. de Freitas, Carolina Q. Sacramento, Lucas Villas Bôas Hoelz, Mônica M. Bastos, Mayara Mattos, Erick Correia Loiola, Pablo Trindade, Yasmine Rangel Vieira, Giselle Barbosa-Lima, Hugo C. de Castro Faria Neto, Nubia Boechat, Stevens K. Rehen, Karin Brüning, Fernando A. Bozza, Patrícia T. Bozza, Thiago Moreno L. Souza

## Abstract

Chikungunya virus (CHIKV) causes a febrile disease associated with chronic arthralgia, which may progress to neurological impairment. Chikungunya fever (CF) is a consolidated public health problem, in tropical and subtropical regions of the world, where control of CHIKV vector, mosquitos of the *Aedes* genus, failed. Since there is no vaccine or specific treatment against CHIKV, infected patients receive only palliative care to alleviate pain and arthralgia. Thus, drug repurposing is necessary to identify antivirals against CHIKV. Recently, the structure and activity of CHIKV RNA polymerase was partially resolved, revealing similar aspects with the enzyme counterparner on other positive sense RNA viruses, such as members of the Flaviviridae family. We then evaluated if sofosbuvir, clinically approved against hepatitis C virus RNA polymerase, which also aims to dengue, Zika and yellow fever viruses replication, would inhibit CHIKV replication. Indeed, sofosbuvir was 5-times more selective in inhibiting CHIKV production in human hepatoma cells than ribavirin, a pan-antiviral drug. Although CHIKV replication in human induced pluripotent stem cell (iPS)-derived astrocytes was less sensitive to sofosbuvir’s, compared to hepatoma cells – this drug still impaired virus production and cell death in a MOI-dependent manner. Sofosbuvir also exhibited antiviral activity *in vivo*, by preventing CHIKV-induced paw oedeme in adult mice, at 20 mg/kg/day, and mortality on neonate mice model, at 40 and 80 mg/kg/day. Our data demonstrates that a prototypic alphavirus, CHIKV, is also susceptible to sofosbuvir. Since this is a clinically approved drug, it could pave the way to become a therapeutic option against CF.

## 1. Introduction

Chikungunya virus (CHIKV) is a member of the *Togaviridae* family, genus alphavirus, which causes febrile debilitating illness associated with arthralgia and skin rash (1). Although prolonged and debilitating joint pain and oedema differentiate CHIKV infection among contemporary arboviruses, like dengue (DENV) and Zika (ZIKV) viruses, most often these agents display similar clinical signs and symptoms during the early phase of infection (1). Severe outcomes of CHIKV infection, leading to acute and convalescent neurological impairment, have also been described (2, 3).

Chikungunya fever (CF) is a consolidated public health problem with substantial impact in tropical and subtropical regions of the world, where *Aedes* spp mosquitoes are prevalent and control measures failed (1). In last 5 years, the Americas, African and Eurasian regions have been severely impacted by CHIKV (4). For instance, in Brazil, since 2014, the Asian and East-Central-South-African (ECSA) genotypes of CHIKV co-circulate (5-7), highlighting a substantial viral activity in a country historically hyperendemic for DENV. Since no specific treatment or vaccine against CHIKV exist, repurposing of clinically approved drugs, preferentially aiming a viral target, is a necessary response against CF.

CHIKV has a positive-sense single-stranded 11.8 kilobase RNA genome, which encodes four non-structural (nsP1-4) and five structural proteins (C, E1, E2, E3 and 6K) (8). Among these proteins, nsP4 encodes for the viral RNA-dependent RNA polymerase (RDRP).Recently, nsP4 structure was partially resolved (9). As other RNA polymerases from positive sense RNA viruses, CHIKV nsP4 possesses well-conserved motifs, such as D-x(4,5)-D and GDD, spatially juxtaposed, wherein Asp binds Mg_2_+ and Asn selects ribonucleotide triphosphates over dNTPs, determining RNA synthesis (10). Moreover, since the RDRP activity is absent on host cells, it constitutes a suitable target for antiviral intervention.

We, and others, have demonstrated that sofosbuvir, a clinically approved against hepatitis C virus (HCV) (11-13), also inhibits the replication of flaviviruses, like ZIKV, DENV and yellow fever virus (YFV) (14-19). Sofosbuvir is a safe and a well-tolerated drug, from 400 to 1200 mg daily in 24 weeks regimen. Sofosbuvir is a uridine monophosphate prodrug that requires the removal of the phosphate protections to enter a pathway that yields sofosbuvir triphosphate, the pharmacological active compound as antiviral (11). Although hepatic cells have the most effective system to remove sofosbuvir’s phosphate protections, functional assays reveals that other cells, relevant for arboviruses infection, may also activate sofosbuvir (11, 16, 20). As expected for a nucleotide analogue, sofosbuvir inhibit the RNA polymerase from different members of the Flaviviridae family, HCV, ZIKV, DENV, YFV (14-19). The conservation of RDRP domain of CHIKV nsP4, when compared to other viral RNA polymerases, led us to hypothesized that CHIKV could also be susceptible to sofosbuvir. Indeed, we originally demonstrated, by cellular assays and animal models, that CHIKV is susceptible to sofosbuvir.

## 2. Results

### 2.1. CHIKV RNA polymerase, nsP4, as the predictive target for sofosbuvir

We considering the homology among viral RDRP to evaluate whether sofosbuvir docks on CHIKV RNA polymerase. For comparisons, the binding mode of sofosbuvir-triphosphate (SFV) and the natural substrate, UTP, were analyzed on the nsP4 model. Three docking simulations for each ligand (totalizing 30 poses per ligand) were carried out. The poses with the lowest energy was selected for analysis (Table 1 and Figure 1). SFV and UTP have similar modes of interaction, but different energy values, respectively, -78.41 and -108.78 arbitrary units (a.u.) (with respect to MolDock scores) (Table 1). Moreover, SFV interacted via H-bonds with Asn348, Ile369, Gly370, Asp371, and Cys411 (H-bond energy = -6.97 a.u.), whereas UTP formed H-bonds with Asn348, Ile369, and Gly370 (bond energy = -3.11 a.u.) (Table 1 and Figure 1). Both SFV and UTP also formed electrostatic attractive interactions with the two Mg^2+^ ions and repulsive interactions with Asp371. Consequently, SFV and UTP displayed electrostatic interaction energies of -117.12 a.u. and -112.84 a.u., respectively (Table 1 and Figure 1). SFV and UTP use similar amino acid residues for steric interactions Phe280, Asn344, Asn348, Ala367, Phe368, Ile369, Asp371, Asp372, Asn373, Ile374, and Cys411, resulting in energies equal to -24.50 a.u and 48.76 a.u, respectively. Nevertheless, minor differences with respect to steric interaction were observed: SFV docks onto Thr345 and Phe410, whereas UTP interacts with Leu250 and Phe251.

**Table 1:** Summary of the interactions of SFV and UTP to nsP4 model of CIKV.

| # | nsP4 model of CHIKV |  |  |  |  |  |  |
| --- | --- | --- | --- | --- | --- | --- | --- |
|  | H-bond energy | Residues (H-bond) | Electrostatic Interaction energy | Residues and Co-Factor (Electrostatic) | Steric interaction energy by PLP<sup>b</sup> | Residues and Co-Factor (Steric) | MolDock score |
| **SFV** | -6.97<sup>a</sup> | Asn348, Ile369, Gly370, Asp371, Cys411 | -117.24 | Asp371, Mg<sup>2+</sup> | -24.50 | Phe280, Asn344, Thr345, Asn348, Ala367, Phe368, Ile369, Gly370, Asp371, Asp372, Asn373, Ile374, Phe410, Cys411 | -78.41 |
| **UTP** | -3.11 | Asn348, Ile369, Gly370 | -112.84 | Asp371, Mg<sup>2+</sup> | -48.76 | Leu250, Phe251, Phe280, Asn344, Thr345, Asn348, Ala367, Ile369, Gly370, Asp371, Asp372, Asn373, Ile374, Cys411 | -108.78 |

<sup>a</sup> Arbitrary units; <sup>b</sup>Piecewise linear potential(21).

**Figure 1.**
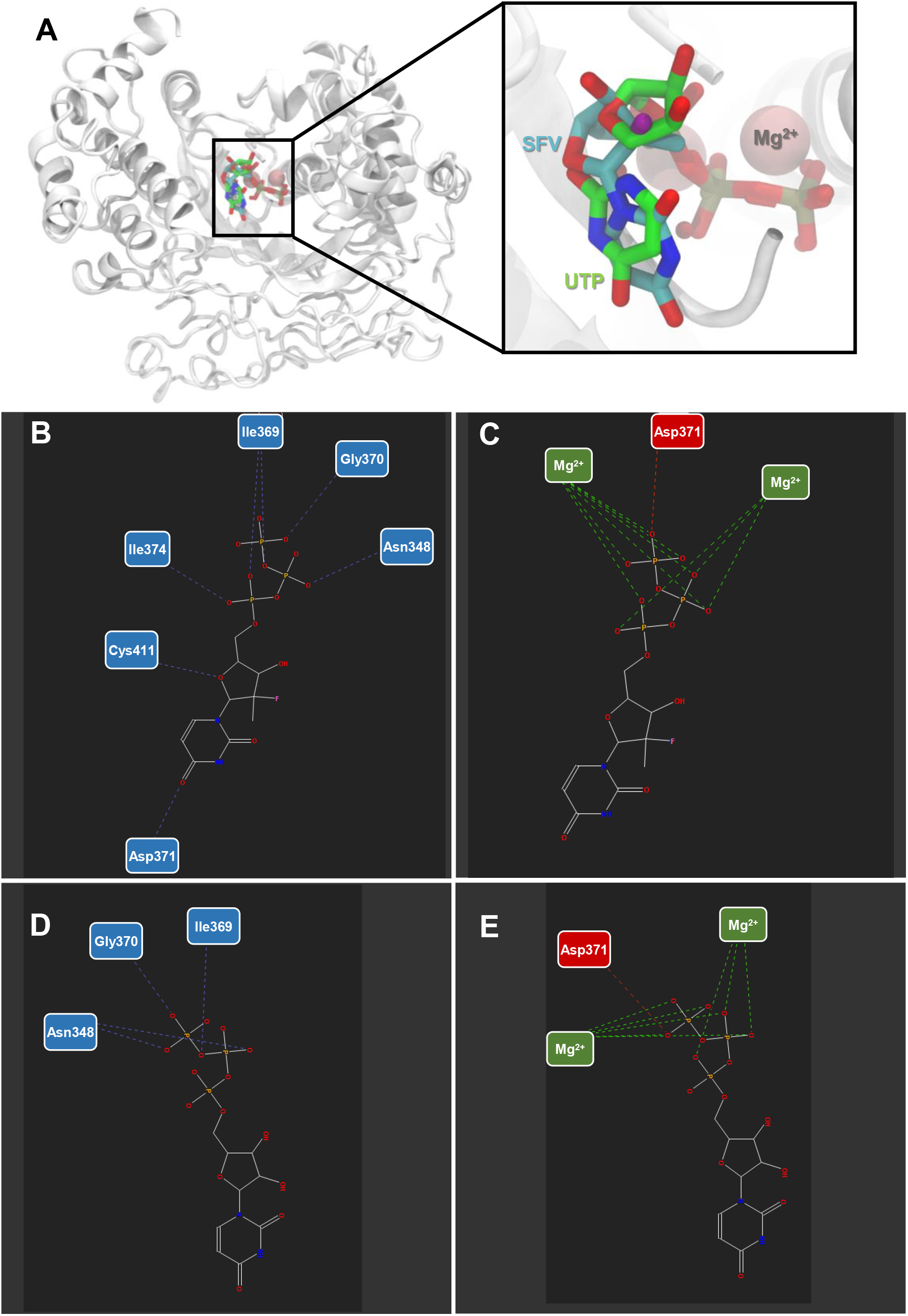
Sofosbuvir triphosphate and nsP4 interactions. (A) Structural representation of the nsP4 model of CHIKV and its interaction with SFV and UTP. Hydrogen bonds and electrostatic interactions between (B; C) SVF and (D; F) UTP, and nsP4 model of CHIKV. The interactions are represented by blue (H-bonds), green (attractive electrostatic interactions), and red (repulsive electrostatic interactions) interrupted lines. The nitrogen atoms are shown in blue, oxygen in red, fluor in pink, and the carbon chain in gray.

### 2.2. CHIKV is susceptible to sofosbuvir *in vitro*

To evaluate if sofosbuvir is indeed endowed with anti-CHIKV activity, phenotypic antiviral assays were performed in human cells previously associated to peripheral virus replication and invasion of the nervous system (22), respectively, hepatoma (Huh-7) and astrocytes derived from induced pluripotent stem (iPS) cells. Supernatants from these infected cultures were harvested and tittered in Vero cells. We observed a dose-dependent inhibition of CHIKV production in hepatoma cells (Figure 2 and Table 2), which is known to possess the machinery to convert the sofosbuvir prodrug to the pharmacologically active metabolite (12). Sofosbuvir was two-fold more potent and 25 % less cytotoxic than ribavirin (Table 2). Consequently, selectivity index (SI) for sofosbuvir was almost five-fold better than for ribavirin, a pan-antiviral drug. Moreover, astrocytes succumb to CHIKV infection in a multiplicity of infection (MOI)-dependent manner, and sofosbuvir partially prevented cell mortality (Figure 3A). Accordingly, sofosbuvir decreased CHIKV replication on astrocytes by 50 % at MOI of 1 (Figure 3B).

**Figure 2.**
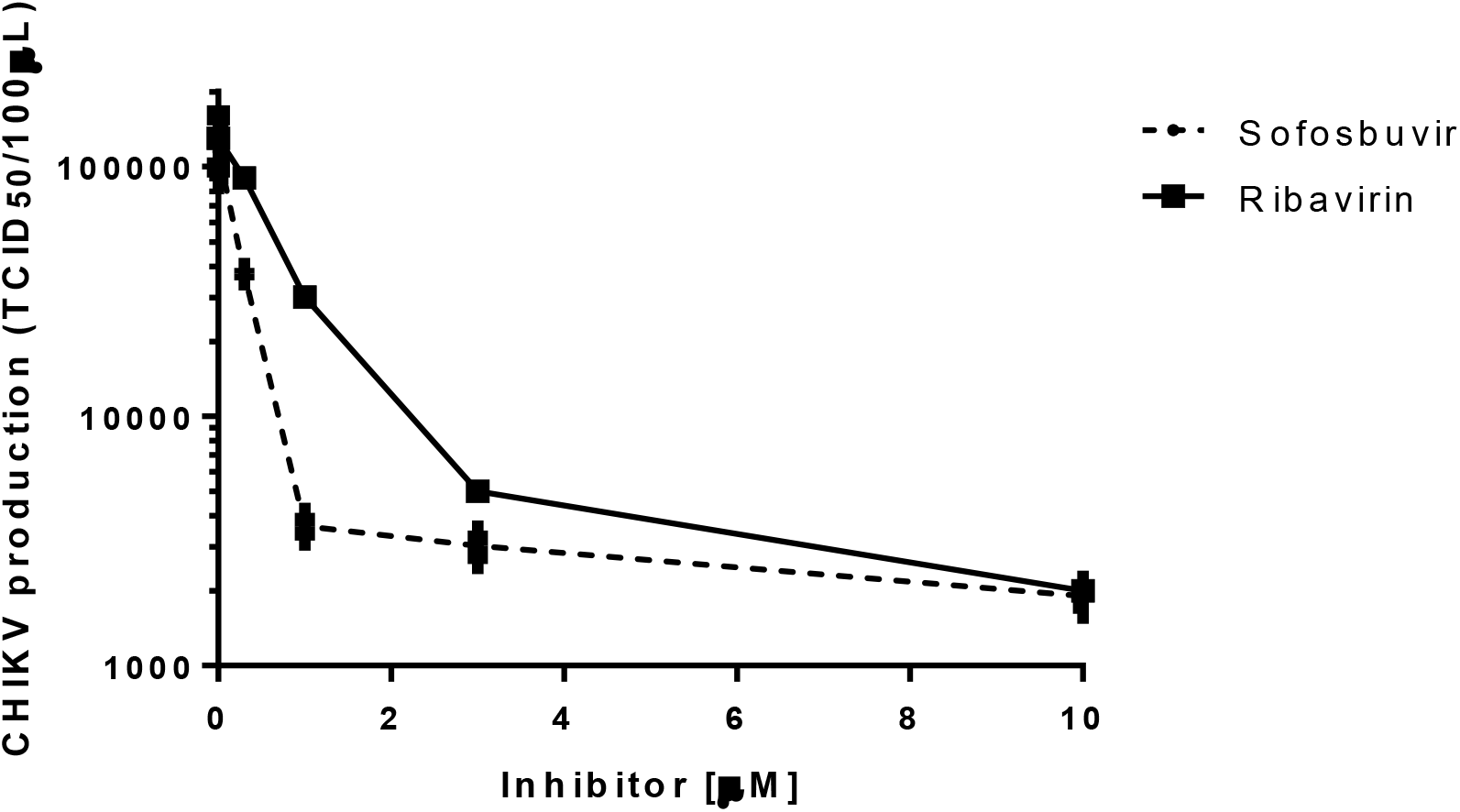
Pharmacology of Sofosbuvir against CHIKV. Huh-7 were infected with CHIKV at MOI of 0.1 and exposed to various concentrations of sofosbuvir or ribavirin for 24 h. Supernatant was harvested and tittered in Vero cells by TCID_50_/mL. The data represent means ± SEM of three independent experiments.

**Table 2.** Pharmacological parameters associated to drug inhibition of CHIKV replication.

| Drug | EC<sub>50</sub> (µM) | CC<sub>50</sub> (µM) | SI* |
| --- | --- | --- | --- |
| Sofosbuvir | 0.8 ± 0.08 | 402 ± 32 | 502 |
| Ribavirin | 1.9 ± 0.3 | 298 ± 22 | 157 |

\*SI – Selectivity Index (CC<sub>50</sub>/EC<sub>50</sub>)

**Figure 3.**
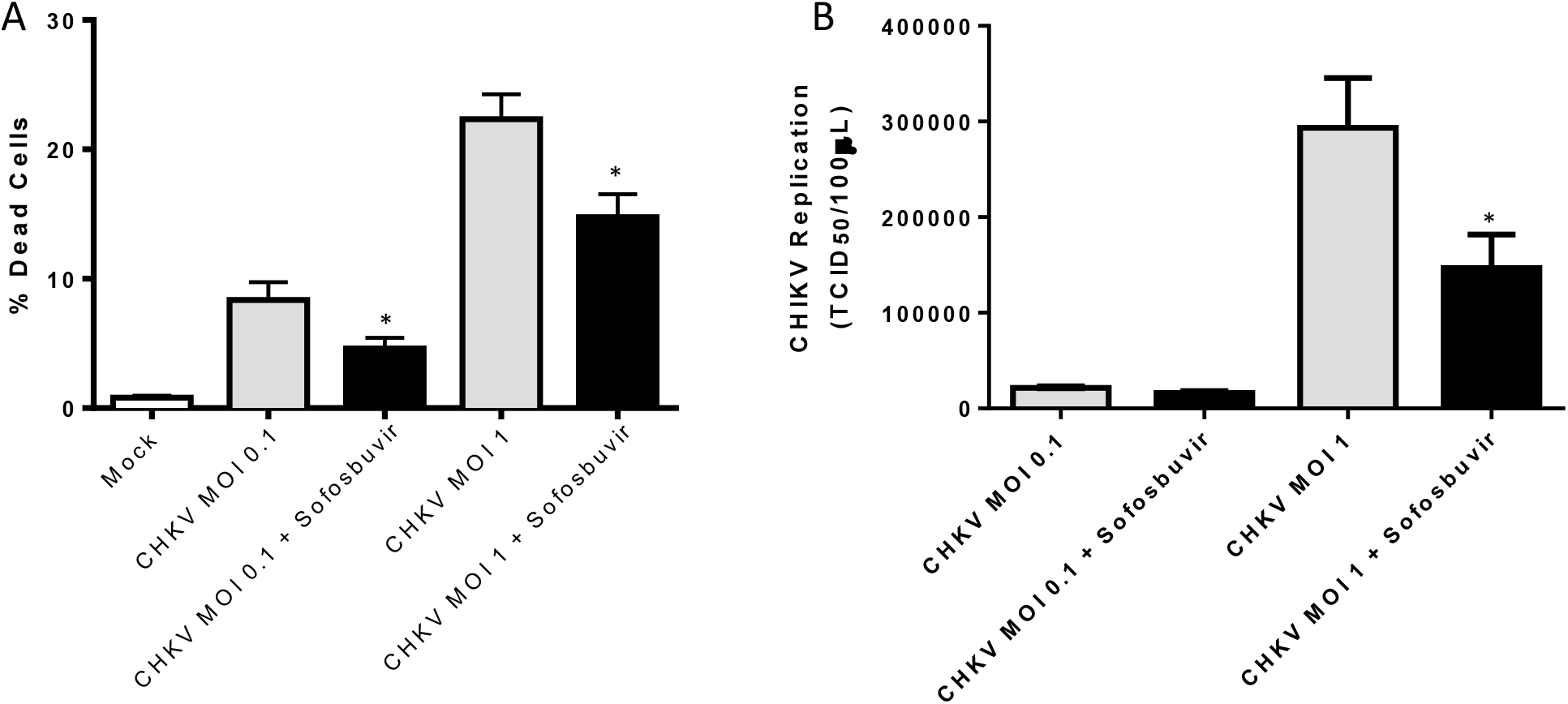
Sofosbuvir inhibits CHIKV replication in human iPS cell-derived astrocytes. Astrocytes were infected at the indicated MOIs and treated with sofosbuvir at 10 µM. After 5 days, cells were labeled for activated caspase-3/7 and propidium iodide (A) and virus in the supernatant tittered in Vero cells (B). The data represent means ± SEM of three independent experiments performed with five technical replicates. \**P <* 0.05 for the comparison between the infected untreated (gray bars) and treated (black bars) groups.

### Sofosbuvir protected CHIKV-infected mice, using models of arthralgia and severe infection

To analyze whether *in vitro* results translate into a systemic protection, we treated CHIKV-infected mouse with sofosbuvir. Initially, treatment was performed in the arthralgia mouse model, in which sofosbuvir was given orally (20 mg/kg/day) one-hour prior to the injection of 2×10^4^ TCID_50_ into the right rind paw. We observed that early after infection paw oedema did not ameliorate with sofosbuvir, suggesting that this drug had now effect on the swelling associated with the insult caused by the injection (Figure 4A-C). Importantly, in the untreated and infected mice, paw oedema was more intense in the following days, whereas the treated animals displayed no differences to mock-infected mice (Figure 4D-F and Figure 5).

**Figure 4.**
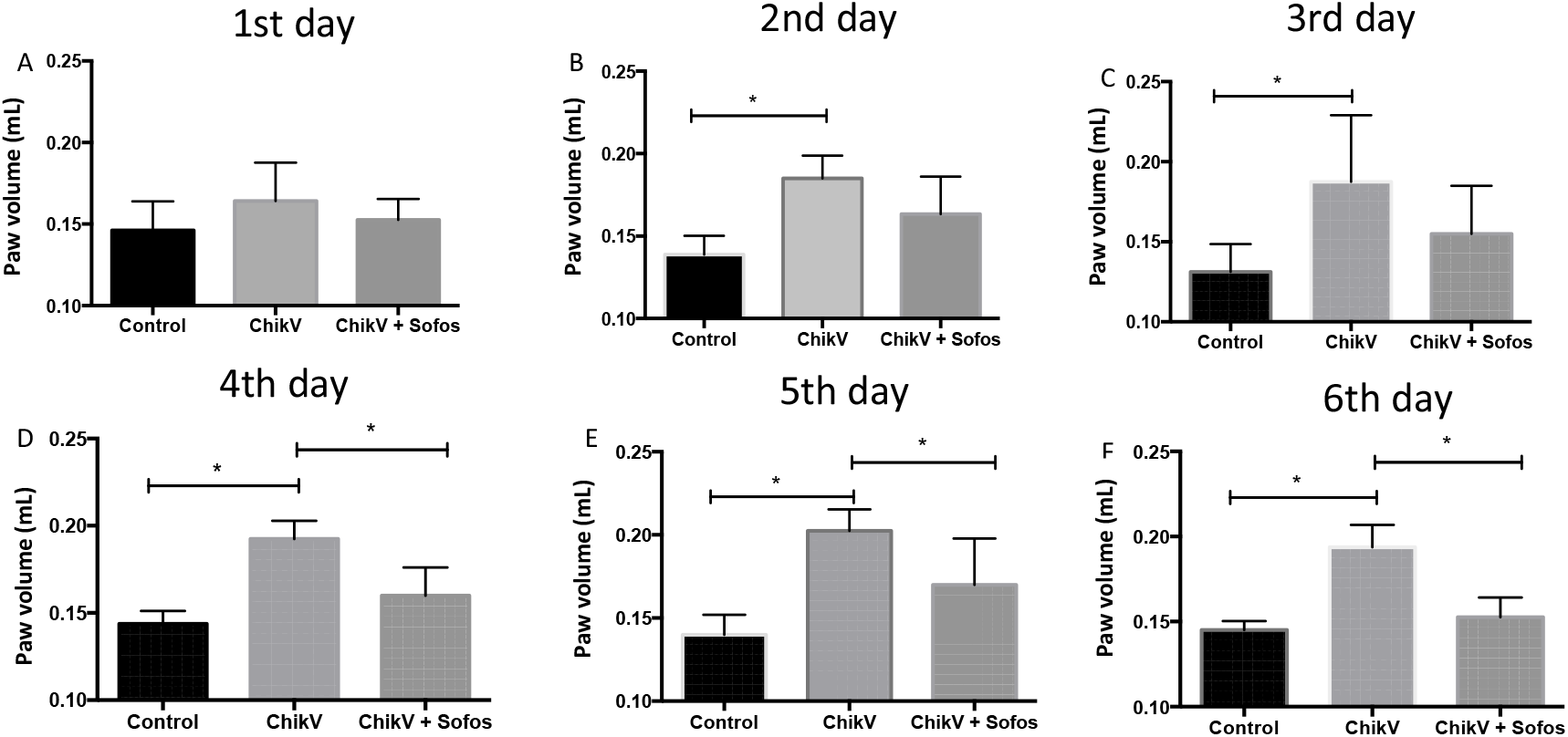
Sofosbuvir ameliorates CHIKV-induced paw oedema. Male Swiss Webster mice (20-25 g) received RPMI medium (Control) or 10^4^ TCID_50_ of CHIKV into 50 μl per paw in the ventral side of the right hind foot. Oral treatment with sofosbuvir (sofos) (20 mg/kg/day) started one hour prior to infection. Panels A to F indicates the days after infection, from the first to the sixth, when paw volume was measured in the hydropletismometer. Paw oedema is interpreted by the increase in paw volume over control in the *p<0.05 Tukey’s multiple comparisons test (n=8/group).

**Figure 5.**
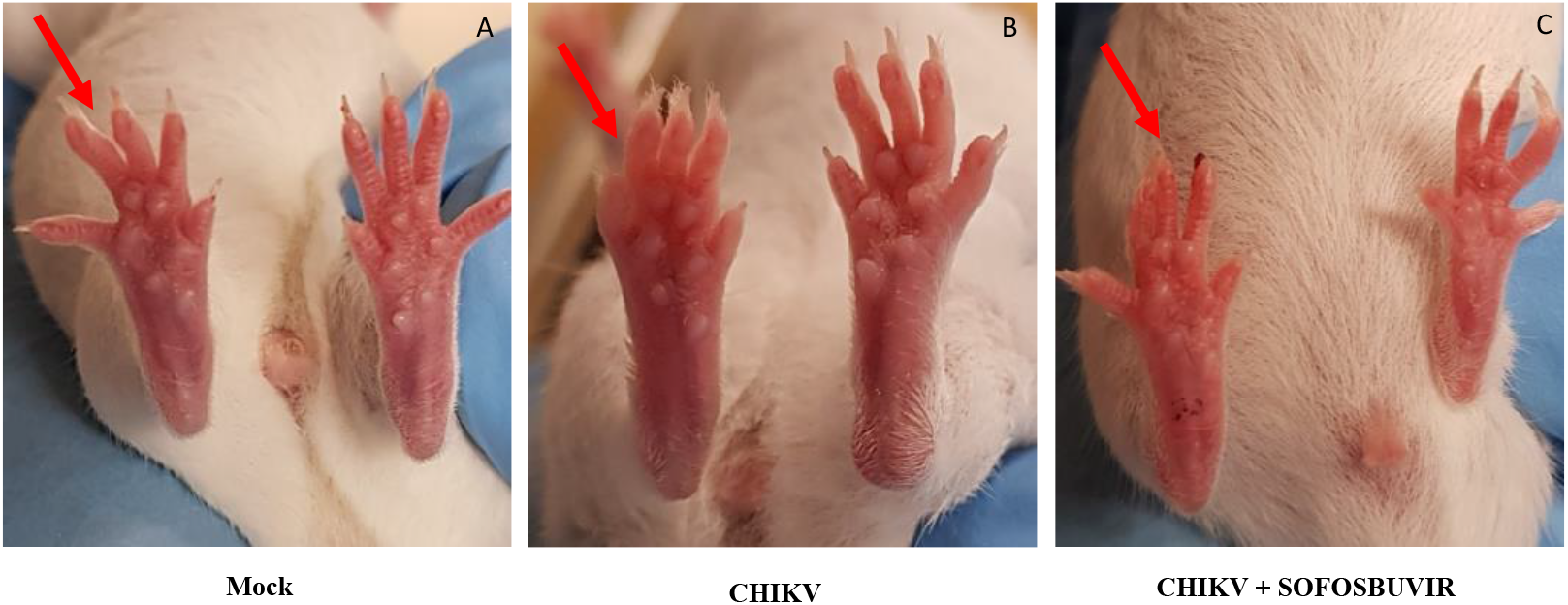
Representative CHIKV-associated paw oedema and sofosbuvir’s antiviral effect on the 6^th^ day after infection. (A) Control, (B) CHIKV and (C) CHIKV + sofosbuvir. Arrows indicate the infected paw.

Subsequently, we studied sofosbuvir’s ability to enhance the survival of CHIKV-infected neonatal mice. Three days-old Swiss mice were infected with CHIKV (2×10^5^ TCID_50_) intraperitoneally. Treatment was carried out daily, initially with 20 mg/kg, also by intraperitoneal injection, beginning at one day prior to infection (pre-treatment) or on second day after infection (late-treatment). Although pre-treatment doubled the mean time of survival (T_50_) when compared to mock-infected animals, all infected mice deceased after 6 days of infection (Figure 6A). Late-treatment had marginal contribution to enhance the T_50_ of mice survival (Figure 6A). Of note, post-natal development of the infected mice varied only marginally among the groups (Figure 6B). Under the same experimental conditions of infection, we next performed pre-treatments with different doses of sofosbuvir. Doses of 40 and 80 mg/kg/day doubled and tripled the percentage of animal survival (Figure 6C). At 80 mg/kg/day animals post-natal development were significantly superior to the infected controls (Figure 6D).

**Figure 6.**
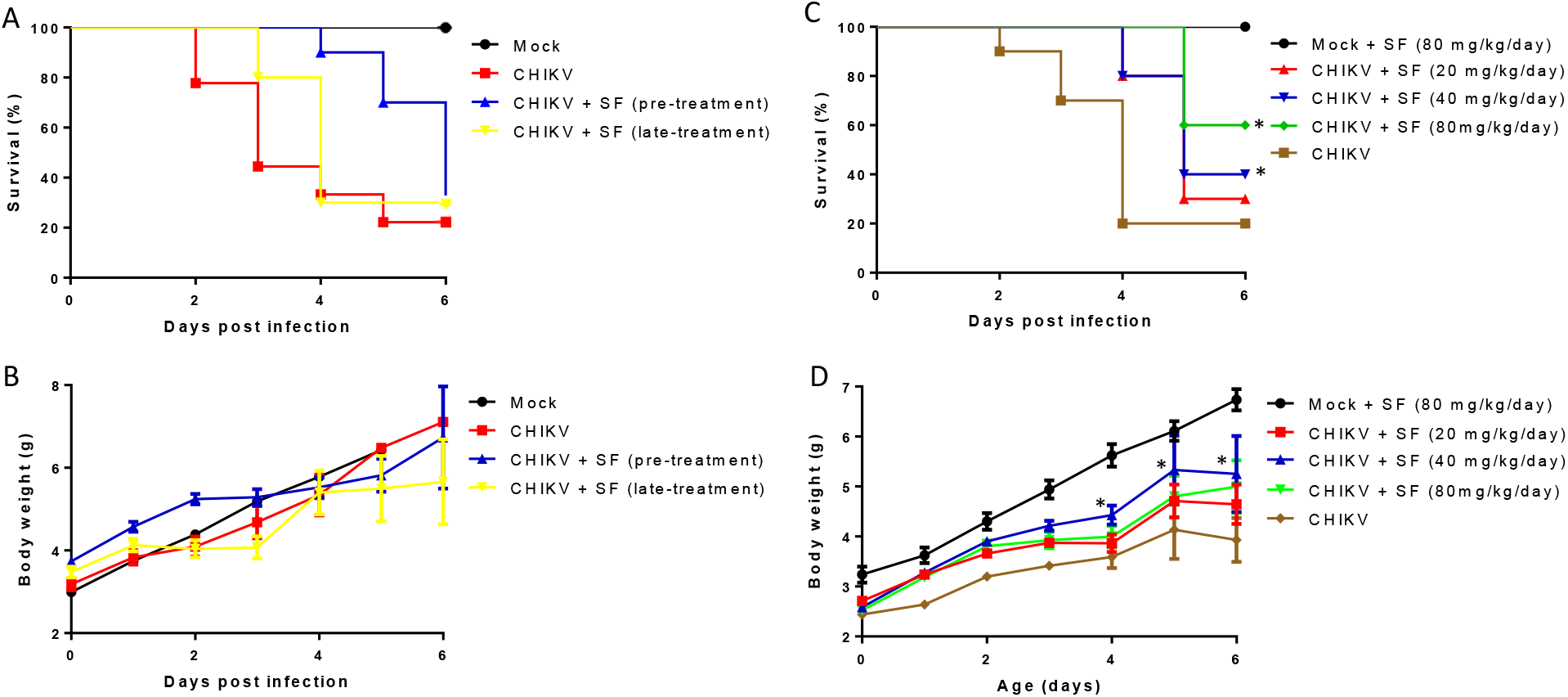
Sofosbuvir, at concentrations of 40 and 80 mg/kg/day, increases survival and inhibits weight loss of CHIKV-infected mice. Three-day-old Swiss mice were infected with CHIKV (2×10^5^ TCID_50_) and treated with sofosbuvir (SF) either 1 day before (pre-treatment) or 2 days after infection (late-treatment). Survival (A and C) and weight variation (B and D) were assessed during the course of treatment. Panels A and B represent experiments of both pre- and late-treatment with sofosbuvir at 20 mg/kg/day. Panels C and D represent pre-treatment with indicated concentrations of sofosbuvir. Survival was statistically assessed by Log-rank (Mentel-Cox) test. Differences in weight are displayed as the means ± SEM, and two-way ANOVA for each day was used to assess the significance. Independent experiments were performed with at least 10 mice/group (n = 30). * P < 0.01.

Since some infected and untreated animals survived from experiments described in Figures 6, we evaluated if they had neuromotor sequelae and compared to treated survivors. Animals were held in a supine position with all four paws facing up, and then released. The time to flip over onto its stomach with all four paws touching the surface was measured, as proxy of neuromotor function. CHIKV-infected mice took a median time of 10-20 second to get to the upright position, whereas mock-infected animals did it immediately (Figure 7). Importantly, CHIKV-infected and sofosbuvir pre-treated animals did not present neuromotor sequela, meaning that these animals are healthier than infected controls (Figure 7). Of note, although the late-treatment diminished the median time associated with neuromotor sequela, some animals displayed a behaviour similar to CHIKV-infected animals, making these groups statistically indistinguishable. Altogether, our data suggests that sofosbuvir also inhibits CHIKV replication *in vivo*, ameliorating animals’ arthralgia, enhancing survival and preserving neuromotor function.

**Figure 7.**
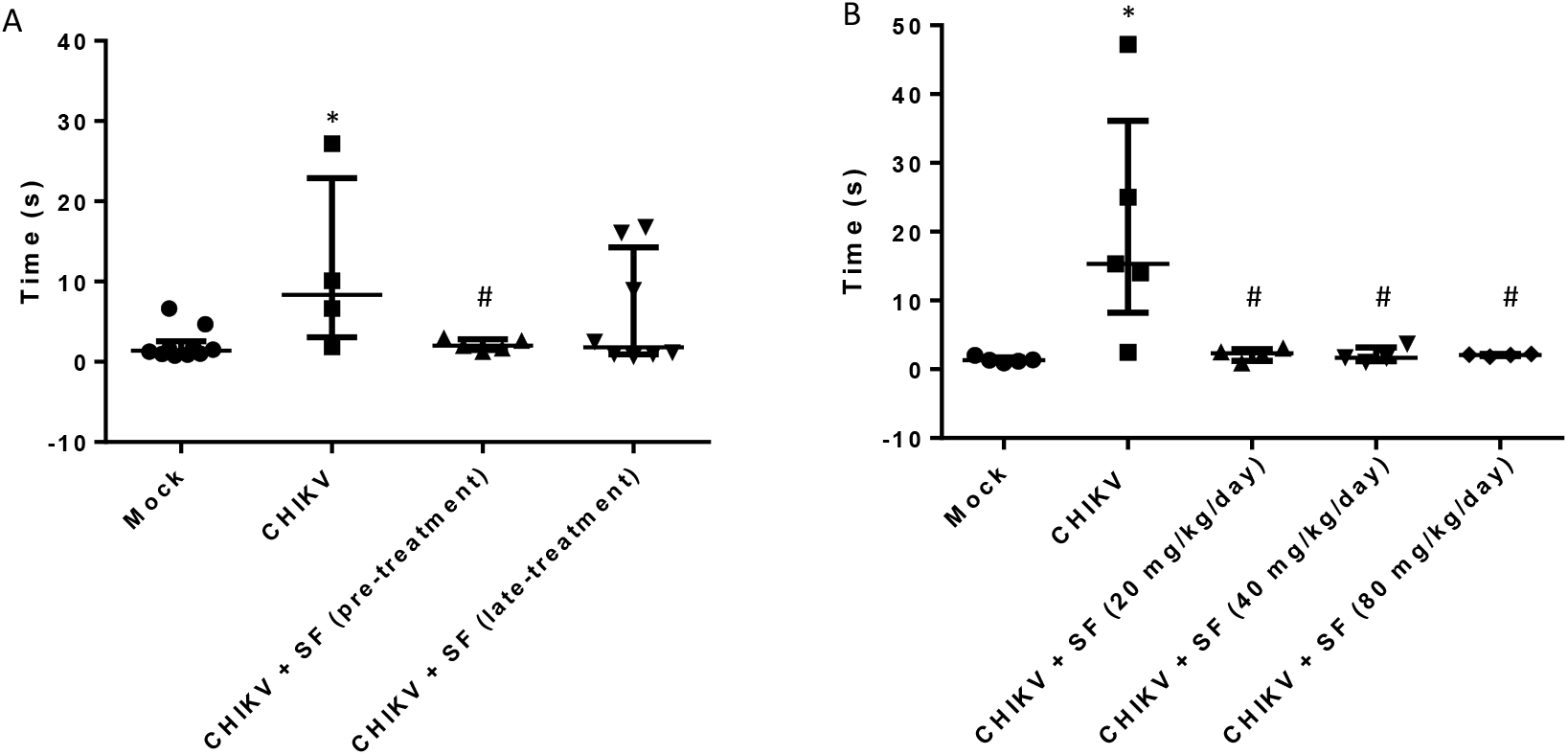
Sofosbuvir prevents neuromotor impairment in CHIKV-infected mice. Three-day-old Swiss mice were infected with CHIKV (2 × 10^5^ TCID_50_) and treated with sofosbuvir (SF) beginning 1 day before infection (pre-treatment) or on the 2^nd^ day after infection (late-treatment). (A) Treatment was performed with 20 mg/kg/day. (B) Pre-treatment was performed with indicated concentrations. At the sixth day after infection, animals were turned backwards and allowed up to 60 s to return to the upright position. The results are presented as the means ± SEM. This was a routine measure and at least 10 animals per group were analysed. Student’s t test was used to compare untreated CHIKV-infected mice with other groups individually. * P < 0.01 mock- vs CHIKV-infected animals. # P < 0.01 untreated vs treated animals.

## 3. Discussion

CHIKV is among the reemergent arboviruses in the early 21^st^ century. Although firstly characterized in the 1950’s in Tanzania (23), since the 2000’s CHIKV activity increased worldwide and reached the new world (24). CF is estimated to cause disability-adjusted life years (DALYs) lost around 45.26 per million people (25). In Brazil, CHIKV was introduced during 2014 (6, 7), when Asian genotype was confirmed in the North Region of Brazil (Oiapoque, Amapá state) and ECSA genotype was identified in the Northeastern region of Brazil (Feira de Santana, Bahia state). The ECSA genotype was subsequently detected throughout Brazil. To the best of our knowledge, Brazil is a rarely case where two genotypes of CHIKV co-circulate. Since early 2016 (summer in the southern hemisphere), DENV, ZIKV and CHIKV co-circulate in Americas, and CHIKV became the most prevalent arbovirus in Brazilian overcrowded cities, like Rio de Janeiro (26). Due to absence of vaccine and specific antiviral treatment, CHIKV prevention depends basically on vector control, whereas patients with CF receive palliative care with nonsteroidal anti-inflammatory drugs (NSAIDs) or corticoids depending on phase of the disease (27).

CHIKV possesses a complex and not fully understood pathogenesis, virus replicate in peripheral organs and may invade the nervous system and synovial fluid (22, 28). Recent efforts to identify substances against CHIKV were carried out, leading to the discovery of chloroquine (29), berberine, abamectin and ivermectin (30). Among these substances, ivermectin was shown to inhibit Flaviviruses NS3 helicase activity (31). By analogy, this drug may also target a CHIKV protein, like nsp2 – which seems to possesses an helicase activity (32). The other identified compounds, such as the alkaloid berberine, target cellular rather than viral pathways (33). We, and others, have shown that sofosbuvir is endowed with antiviral activity against flaviviruses (15-17). The recent advances in the CHIKV nsP4 RDRP core domains structure and function highlighted to the presence of conserved motifs among RNA polymerases from positive-sense RNA viruses (9). Indeed, sofosbuvir docked onto CHIKV nsP4 using conserved amino acid residues, also required for binding of UTP, the natural substrate. Sofosbuvir inhibited CHIKV replication in human hepatoma cells. These cells were used because they represent one of the most efficient *in vitro* models to convert sofosbuvir to the active metabolite and liver is a relevant organ for CHIKV pathogenesis (22, 28). Growing evidence indicates that CHIKV may impair neurological function, by directly invading the nervous system (28, 34, 35). We used a sophisticated human iPS-derived astrocyte culture to show that sofosbuvir inhibits CHIKV replication and virus-associated cell mortality, although in lesser effective manner when compared to huh-7 cells. Accordingly, in the neonatal mouse model of CHIKV infection, sofosbuvir enhanced animal survival at doses higher than required to produce the same effect on ZIKV-infected pups (16). Consistently, sofosbuvir is safe to be used clinically up to 1200 mg per day (12). With respect to the time frame of opportunity to treat CHIKV-infected mice, a narrow window was observed – similarly to what we have noticed towards ZIKV (36). In other acute virus infections, like influenza, mortality is dramatically reduced when neuraminidase inhibitors are administered early in the time course of infection, such as within 2.5 days of infection (37). The identification of individuals at higher risk, to receive sofosbuvir prophylactically, or very early after infection represent one of the challenges to translate our date into public health intervention.

Moreover, we observed that, at experimental infection conditions milder than required for animal mortality, sofosbuvir, at reference dose for pre-clinical studies, 20 mg/kg/day (13), protected arthralgia-related paw oedema. Thus, it is important to further study whether sofosbuvir could act synergistically with anti-inflammatory drugs to improve the quality of life for patients with CHIKV-associated chronic arthralgia. Patients with CF very often present arthralgia and impairment of the neuromuscular function, causing a debilitating condition which contributes to the burden of disease (28). Consistently, using survivor animals, we observed that sofosbuvir protected CHIKV-infected mice form neuromotor sequalae when compared to untreated animals. Under this behaviour test, it is likely that both direct neuromotor sequelae and/or problems on mice articulations may contributed to the high time required for CHIKV-infected animals to turn from back to upright position.

Altogether, our data reveals that CHIKV is susceptible to sofosbuvir, highlighting that other genetically distinct and clinically important viruses phylogenetically distributed among members of the Togaviridae and Flaviviridae families could also be susceptible to this drug. Wider use of sofosbuvir, beyond HCV, may represent a safer antiviral option than ribavirin. Finally, in the context of this study, our findings motivate phase II clinical investigations on the new use of sofosbuvir as treatment against CHIKV.

## 4. Material and Methods

### Reagents

The antiviral sofosbuvir (β-d-2’-deoxy-2’-α-fluoro-2’-β-C-methyluridine) was donated by the BMK Consortium: Blanver Farmoquímica Ltda; Microbiológica Química e Farmacêutica Ltda; Karin Bruning & Cia. Ltda, (Taboão da Serra, São Paulo, Brazil). Ribavirin was received as a donation from the Instituto de Tecnologia de Farmacos (Farmanguinhos, Fiocruz). All small molecule inhibitors were dissolved in 100 % dimethylsulfoxide (DMSO) and subsequently diluted at least 10^4^-fold in culture or reaction medium before each assay. The final DMSO concentrations showed no cytotoxicity. The materials for cell culture were purchased from Thermo Scientific Life Sciences (Grand Island, NY), unless otherwise mentioned.

### Cells

African green monkey kidney (Vero) and human hepatoma (Huh-7) cells were cultured in DMEM. The culture medium of each cell type was supplemented with 10 % fetal bovine serum (FBS; HyClone, Logan, Utah), 100 U/mL penicillin, and 100 µg/mL streptomycin(38, 39) at 37 °C in 5 % CO_2_.

### Virus

CHIKV (Asian strain) was donated by Dr. Amilcar Tanuri. CHIKV was propagated in Vero cells at a multiplicity of infection (MOI) of 0.1. Infection was carried out for 1 h at 37 °C. Next, the residual virus particles were removed by washing with phosphate-buffered saline (PBS), and the cells were cultured for an additional 2 to 5 days. After each period, the cells were lysed by freezing and thawing and centrifuged at 1,500 × *g* at 4 °C for 20 min to remove cellular debris. Virus titters were determined by classical 10-fold dilution and tissue cytopathic infectious dose 50 (TCID_50_)/mL calculation.

### Cytotoxicity assay

Monolayers of cells 2 to 5 × 10^4^ cells/well in 96-well plates were treated for 5 days with various concentrations of sofosbuvir or ribavirin as a control. Then, 5 mg/ml 2,3-bis-(2-methoxy-4-nitro-5-sulfophenyl)-2*H*-tetrazolium-5-carboxanilide (XTT) in DMEM was added to the cells in the presence of 0.01 % of N-methyl dibenzopyrazine methyl sulfate (PMS). After incubating for 4 h at 37 °C, the plates were read in a spectrophotometer at 492 nm and 620 nm (40). The 50 % cytotoxic concentration (CC_50_) was calculated by a non-linear regression analysis of the dose–response curves.

### Yield-reduction assay

Monolayers of 5 × 10^4^ Huh-7 cells/well in 96-well plates were infected with CHIKV at the MOI of 0.1 for 1 h at 37 °C. The cells were washed with PBS to remove residual viruses, and various concentrations of sofosbuvir, or ribavirin as a positive control, in DMEM with 1 % FBS were added. After 24 h, the cells were lysed, the cellular debris was cleared by centrifugation, and the virus titers in the supernatant were determined in Vero cells as TCID_50_/mL. A non-linear regression analysis of the dose-response curves was performed to calculate the concentration at which each drug inhibited the plaque-forming activity of CHIKV by 50 % (EC_50_).

### Generation of human iPSC-derived astrocytes lines

Astrocytes were differentiated from neural stem cells (NSC), 20 × 10³ cells/well in a 96-well plate, obtained from human iPSC of three control cell lines from healthy subjects (41). These cell lines were previously used in other studies from our research group (16). Three cell lines from healthy subjects were obtained from a female subject (GM23279A, available at Coriell Institute - coriell.org) and the other two from male subjects from cells reprogrammed at the D’Or Institute for Research and Education (CF1 & CF2). NSCs were differentiated into astrocytes as described in Yan, 2013 (42). Briefly, NSCs were cultured in differentiation media (1% N2 supplement and 1% FBS in DMEM/F12) for 21 days with media changes every other day and passages every week. After this period, glial cells were grown for 5 weeks in 10% FBS in DMEM/F12 with media changes twice a week prior to use. Cells were infected at MOIs of either 1.0 or 10 for 2 h at 37 °C. Next, the cells were washed, and fresh medium containing sofosbuvir was added. The cells were treated daily with sofosbuvir at the indicated concentrations. Virus titers were determined from the culture supernatant. Cell death was measured by adding 2 µM CellEvent caspase-3/7 reagent and the fluorescent dye ethidium homodimer (43), when the culture supernatants were collected, on the 5^th^ day after infection. Images were acquired with an Operetta high-content imaging system with a 20x objective and high numerical apertures (NA) (PerkinElmer, USA). The data were analyzed using the high-content image analysis software Harmony 5.1 (PerkinElmer, USA). Seven independent fields were evaluated from triplicate wells per experimental condition.

### 3D Modeling of the Chikungunya Virus Nonstructural Protein 4

The amino acid sequence of the nonstructural protein 4 (NSP4) of chikungunya virus (CHIKV, UniProtKB ID: F2YI10) was obtained from ExPASy server (44). The region between Met1-Lys516, part of the nsP4 sequence that includes the whole catalytic core was considered to construct the model using I-TASSER server (45). The I-TASSER methodology is very accurate for the construction of protein models when the sequence identity between the target sequence and the template protein drops below 30 %, where lack of a high-quality structure match may provide substantial alignment errors and, consequently, poor quality models (45, 46). Thus, the final model was validated using two programs: PROCHECK (47) and VERIFY3D (48). PROCHECK analyzes the stereochemical quality and VERIFY3D compatibility analysis between the 3D model and its own amino acid sequence, by assigning a structural class based on its location and environment, and by comparing the results with those of crystal structures with good resolutions (47, 48).

### Molecular Docking

The structures of SFV and UTP were built in the Spartan’14 software (Wavefunction, Inc., Irvine, CA). The docking of the two ligands to the nsP4 model was performed using Molegro Virtual Docker 6.0 (MVD) program (CLC bio, Aarhus, Denmark) (21), which uses a heuristic search algorithm that combines differential evolution with a cavity prediction algorithm (21). The MolDock scoring function used is based on a modified piecewise linear potential (PLP) with new hydrogen bonding and electrostatic terms included. The full description of the algorithm and its reliability compared to other common docking algorithm have been described (21). The two Mg^2+^ ions were set as the center of searching space with a radius value of 10 Å. In addition, the search algorithm MolDock optimizer was used with a minimum of 100 runs and the parameter settings were population size = 500; maximum iteration = 2000; scaling factor = 0.50; offspring scheme = scheme 1; termination scheme = variance-based; crossover rate = 0.90. Due to the stochastic nature of algorithm search, three independent simulations per ligand were performed to predict the binding mode. Consequently, the complexes with the lowest interaction energy were evaluated. The interactions between nsP4 model and each ligand were analyzed using the ligand map algorithm, a standard algorithm in MVD program (21). The usual threshold values for hydrogen bonds and steric interactions were used.

All figures of nsP4 modeling and molecular docking results were edited using Visual Molecular Dynamics 1.9.3 (VMD) program (available for download at http://www.ks.uiuc.edu/Research/vmd/vmd-1.9.3/) (49).

### Animals

Swiss albino mice (*Mus musculus*) (pathogen-free) from the Oswaldo Cruz Foundation breeding unit (Instituto de Ciência e Tecnologia em Biomodelos (ICTB)/Fiocruz) were used for these studies. The animals were kept at a constant temperature (25°C) with free access to chow and water in a 12-h light/dark cycle. The experimental laboratory received pregnant mice (at approximately the 14th gestational day) from the breeding unit. Pregnant mice were observed daily until delivery to accurately determine the postnatal day. We established a litter size of 10 animals for all experimental replicates.

The Animal Welfare Committee of the Oswaldo Cruz Foundation (CEUA/FIOCRUZ) approved and covered (license numbers L-016/2016 and CEUA L-002/2018) the experiments in this study. The procedures described in this study were in accordance with the local guidelines and guidelines published in the National Institutes of Health Guide for the Care and Use of Laboratory Animals. The study is reported in accordance with the ARRIVE guidelines for reporting experiments involving animals(50). If necessary to alleviate animal suffering, euthanasia was performed. The criteria were the following: i) differences in weight gain between infected and control groups >50%, ii) ataxia, iii) loss of gait reflex, iv) absence of righting reflex within 60 seconds, and v) separation, with no feeding, of moribund offspring by the female adult mouse.

### Experimental infection and treatment

#### Neonate model

Three-day-old Swiss mice were infected intraperitoneally with 2 × 10^2^ TCID_50_ of virus(51, 52), unless otherwise mentioned. Treatments with sofosbuvir were carried out with sofosbuvir at 20 mg/kg/day intraperitoneally. Treatment started one day prior to infection (pretreatment) or two days after infection (late treatment). In both cases, treatment was conducted for 6 days. For comparisons, mock-infected and mock-treated groups of animals were used as controls. Animals were monitored daily for survival, weight gain, and virus-induced short-term sequelae (righting in up to 60 seconds).

#### Arthralgia model

Arthralgia model was adapted from previous publications (53, 54). Male Swiss Webster mice (8 weeks-old; 20-25 g) were infected with 10^4^ TCID_50_ in right hind paw towards the ankle. Sofosbuvir was given orally (20 mg/kg), beginning one-hour before the first virus injection. Treatment was conducted for 6 days. Control group was injected with 50 μL of RPMI. Paw oedema was evaluated from the first to the sixth day after infection by hydropletismometer for small volumes (Ugo Basile, Milan, Italy) and data was presented as paw volume (mL).

#### Behavioural tests

To test the righting reflex, animals were tested daily during the course of acute infection. Animals were held in a supine position with all four paws facing up in the air for 5 seconds. Then, animals were released, and the time the animal took to flip over onto its stomach with all four paws touching the surface was measured. A maximum of 60 seconds was given for each trial, and animals were tested twice a day with a 5-minute minimum interval between trials. For each animal, the lowest time was plotted in the graph. Animals that failed the test were included in the graph with a time of 60 seconds.

#### Statistical analysis

All assays were performed and codified by one professional. Subsequently, a different professional analyzed the results before the identification of the experimental groups. This approach was used to keep the pharmacological assays blind. All experiments were carried out at least three independent times, including technical replicates in each assay. The dose-response curves used to calculate the EC_50_ and CC_50_ values were generated by Excel for Windows. The dose-response curves used to calculate the IC_50_ values were produced by Prism GraphPad software 5.0. Significance of survival curves was evaluated using the Log-rank (Mantel-Cox) test. The equations to fit the best curve were generated based on R^2^ values ≥ 0.9. ANOVA, followed by Tukey’s post hoc test, tests were also used, with *P* values <.05 considered statistically significant. The statistical analyses specific to each software program used in the bioinformatics analysis are described above.

## AUTHOR CONTRIBUTIONS

Performed the experiments – ACF, PAR, CSdeF, CQS, LVBH, MMB, MM, ECL, PT, YRV, GB-L

Conceptualized the experiments and fund raising – HCCFN, NB, SKR, KB, FAB, PTB, TMLS

Provided critical material – KB

Study organization - TMLS

All authors revised and approved the manuscript.

## Acknowledgements

Funding was provided by National Council for Scientific and Technological Development (CNPq), Ministry of Science, Technology, Information and Communications (grants including, but not limited to, # 465313/2014-0; Ministry of Education/CAPES (# 465313/2014-0); Research Foundation of the State of Rio de Janeiro (FAPERJ) (grants including, but not limited to, # 465313/2014-0 and Oswaldo Cruz Foundation (Fiocruz).

